# Optimizing the Benefits of Mental Practice on Motor Acquisition and Consolidation with Moderate-Intensity Exercise

**DOI:** 10.1101/2022.11.12.516269

**Authors:** Dylan Rannaud Monany, Florent Lebon, Charalambos Papaxanthis

## Abstract

The optimization of mental practice (MP) protocols matters for sport and motor rehabilitation. In this study, we were interested in the benefits of moderate-intensity exercise in MP, given its positive effects on the acquisition and consolidation of motor skills induced by physical practice (PP). Four experimental groups were tested: i) physical practice without exercise (PP-Rest), ii) mental practice without exercise (MP-Rest), iii) mental practice preceded by Exercise (Exe-MP), and iv) mental practice followed by Exercise (MP-Exe). We hypothesized that exercise before MP would further increase speed and accuracy at a finger-sequence task measured right after MP (potentiation of motor acquisition), whereas exercise after MP would further increase speed and accuracy the day after MP (promotion of motor consolidation). Motor performance (movement speed and accuracy) was measured during a sequential finger tapping task before (Pre-Test), immediately after (Post-Test 0h, acquisition), and one day after practice (Post-Test 24h, consolidation). Results suggest that exercise before MP did not additionally improve motor acquisition in comparison to the MP-Rest group (both for accuracy and speed, p’s>0.05). Interestingly, moderate-intensity exercise after MP further increased performance during motor consolidation (speed, p=0.051; accuracy, p=0.028), at the level of the PP-Rest group. This novel finding represents a promising advance in the optimization of mental practice protocols in sport-related and rehabilitation settings.

## Introduction

Motor learning is an important process to develop a profuse and skilled motor repertoire. Following physical practice (PP), movements are performed faster and more accurately (Karni et al., 1998). The positive effects of PP on motor performance can be observed with different timescales, from a single practice session to several days or weeks of practice. Previous studies reported that a few minutes of practice is enough to improve motor performance on various tasks, such as pointing or sequential finger-tapping (Spampinato et al. 2017; Ruffino et al. 2022; Walker et al., 2003). This fast-learning process is defined as *motor acquisition* and is considered the first step in the development of new motor memories. Motor memory consolidation, which is the stabilization or the improvement of motor skills between (off-line learning) or within practice sessions, necessitates extensive training (Krakauer and Shadmehr, 2006; Ruffino et al., 2021; Truong et al., 2022).

It is of interest that additional activities, such as moderate-intensity exercise, can assist physical practice and further improve skill acquisition and consolidation (Roig et al., 2013; Mang et al., 2014; Statton et al., 2015; Thomas et al., 2016). The positive effect of exercise depends on its intensity, i.e., moderate or high, and its timing with respect to the practice session, i.e., before or after. For instance, Statton et al. (2015) pointed out that a single bout of moderate-intensity exercise before PP promotes motor acquisition without any effect on motor consolidation. Interestingly, moderate-intensity exercise after PP promotes consolidation (Thomas et al., 2016). Roig et al. (2013) reported that high-intensity exercise, performed before or after PP, promotes motor consolidation without effect on motor acquisition. The positive effects of moderate-to high-intensity exercise on motor acquisition and consolidation are justified, at least in part, by exercise-induced neuroplasticity phenomena. We could cite the facilitation of long-term-potentiation (LTP)-like plasticity induction within the primary motor cortex during motor acquisition (Mang et al., 2014; Ziemann et al., 2006), as well as the increase of circulating brain-derived neurotrophic factor (BDNF) and neuro-adrenergic system activity, which are both involved in motor consolidation (Bekinschtein et al., 2014; Kuo et al., 2021; Segal et al., 2012; Skriver et al., 2014).

Although motor learning usually relies on PP, alternative forms of practice also exist. Among these, mental practice (MP) based on motor imagery, namely the mental simulation of movements without concomitant motor output, is well documented. Several studies reported positive effects of MP on motor acquisition and consolidation, considering parameters such as movement speed or accuracy (Gentili et al., 2006, 2010; Rannaud Monany et al., 2022a). The positive effects of MP come from the activation of neural structures involved during PP (Hétu et al., 2013; Kilteni et al., 2018), as well as neuronal adaptations that sustain both motor acquisition and consolidation (Ruffino et al., 2017). Indeed, functional imagery and neuromodulation studies, respectively, showed that motor imagery and actual execution share overlapping cortico-subcortical networks and induce congruent modulations of corticospinal excitability (Grosprêtre et al., 2015; Hardwick et al., 2018). Furthermore, other studies reported that MP induces neural changes after a single practice session (Avanzino et al., 2015; Ruffino et al., 2019) and after extended periods of practice (Pascual-Leone et al., 1995).

One could argue that, if MP and PP involve overlapping neural substrates and induce neural changes within these substrates, moderate-intensity exercise could also promote MP efficiency. However, to the best of our knowledge, the effects of such type of exercise on MP have never been explored. The aim of the following experiment was thus to probe the effects of moderate-intensity exercise on motor acquisition and consolidation induced by MP. We hypothesized that a single bout of exercise performed before MP would promote motor acquisition, while exercise after MP would promote motor consolidation.

## Methods

### Participants

Forty healthy right-handed volunteers from the Sport Science faculty of Dijon (France) participated in the study (17 women, mean age: 23 years old, range: 19 - 29). The participants were randomly assigned to one of the four experimental groups: i) physical practice without moderate-intensity exercise (PP-Rest group, n=10, mean age: 22, range 19-29), ii) mental practice without moderate-intensity exercise (MP-Rest group, n=10, mean age: 25, range 21-27), iii) mental practice preceded by moderate-intensity exercise (Exe-MP group, n=10, mean age: 24, range 21-27), or and iv) mental practice followed by moderate-intensity exercise (MP-Exe group, n=10, mean age: 21, range 18-27). No professional musicians nor professional typists were recruited due to the nature of the motor task (finger tapping). The study complied with the standards set by the Declaration of Helsinki (Version 2013, except pre-registration).

Participants of the three MP groups were asked to complete the French version of the Motor Imagery Questionnaire to assess self-estimation of their motor imagery vividness (Loison et al., 2013). For this questionnaire, the minimum score is 8 (low imagery vividness) and the maximum one is 56 (high imagery vividness). In the current study, the average scores suggest average to good motor imagery ability (Guillot et al., 2008) and do not differ between groups (F_2,27_=0.182, p=0.83; MP-Rest: 41 ±4.08; Exe-MP: 42.2 ±7, and MP-Exe: 42.6 ±6.96).

### Modality of moderate-intensity exercise

Participants of the Exe-MP and MP-Exe groups were asked to perform a moderate-intensity cycling exercise (*Wattbike Ltd*) for 30 minutes and to maintain themselves at 65-70% of maximal theoretical cardiac frequency (Formula for maximal theoretical cardiac frequency: 220 – age [Fox et al., 1971]; global mean ±SD at 65-70% of maximal theoretical cardiac frequency: 132.74 ±2.89 beats per minute [Riebe et al., 2018]). Participants were self-monitored with a cardio-frequency meter (*Cyclus 2*) directly connected to the bike. The recorded cardiac frequency was displayed in real-time on the bike’s screen. The experimenter regularly checked the compliance of the participants to the given instructions during the exercise. Resistance of the bike was set to minimal for all the participants.

### Behavioral tasks

The motor task consisted of a computerized version of the sequential finger-tapping task (Rannaud Monany et al., 2022a). The participants were seated in front of a customized keyboard and performed a sequence of six movements with their left-hand fingers in the following order: 1-2-4-5-3-5 (1: thumb, 2: index finger, 3: middle finger, 4: ring finger, 5: pinky), as fast and accurately as possible during thirty-second trials. Before the beginning of the experiment, participants were asked to perform the required sequence in front of the experimenter to ensure they understood the task correctly. During the practice session, participants of the PP group had to physically execute the movement whereas participants of the MP group had to imagine it (see below for details).

### Experimental procedures

The experimental protocol included three test sessions (Pre-Test, Post-Test 0h, and Post-Test 24h) and a practice session. Each test session consisted of two actual trials of thirty seconds during which participants were asked to repeat the six-movements sequence, described above, as fast and accurately as possible. After Pre-Test, all groups (physically or mentally) repeated the movements for a total amount of practice of three hundred seconds. For the PP-Rest group, the practice session consisted of ten 30-second trials with the same instructions as in the Pre-Test. Each actual trial was followed by thirty seconds of rest. For the participants of the MP groups, the practice session consisted of three blocks of ten 10-second imagined trials of the sequence. We chose 10-second trials for MP to minimize the deleterious effects of long trials on motor imagery clarity (Rozand et al., 2015). Each imagined trial was followed by ten seconds of rest. Blocks were separated by 1 minute of rest. To optimize the benefits of MP for a such motor task (Lebon et al., 2018), the participants of the MP groups imagined the motor sequence with kinesthetic modality based on the following instructions: “*try to imagine yourself performing the motor task as fast and accurately as possible, by feeling your fingers moving as if you were moving it*”. Depending on their groups, participants remained at rest (PP-Rest and MP-Rest groups) or performed a bout of moderate-intensity exercise before the Pre-Test session (Exe-MP group) or after the Post-Test 0h session (MP-Exe group). A schematic representation of the experimental procedures is depicted in Figure 1.

**Figure 1:**
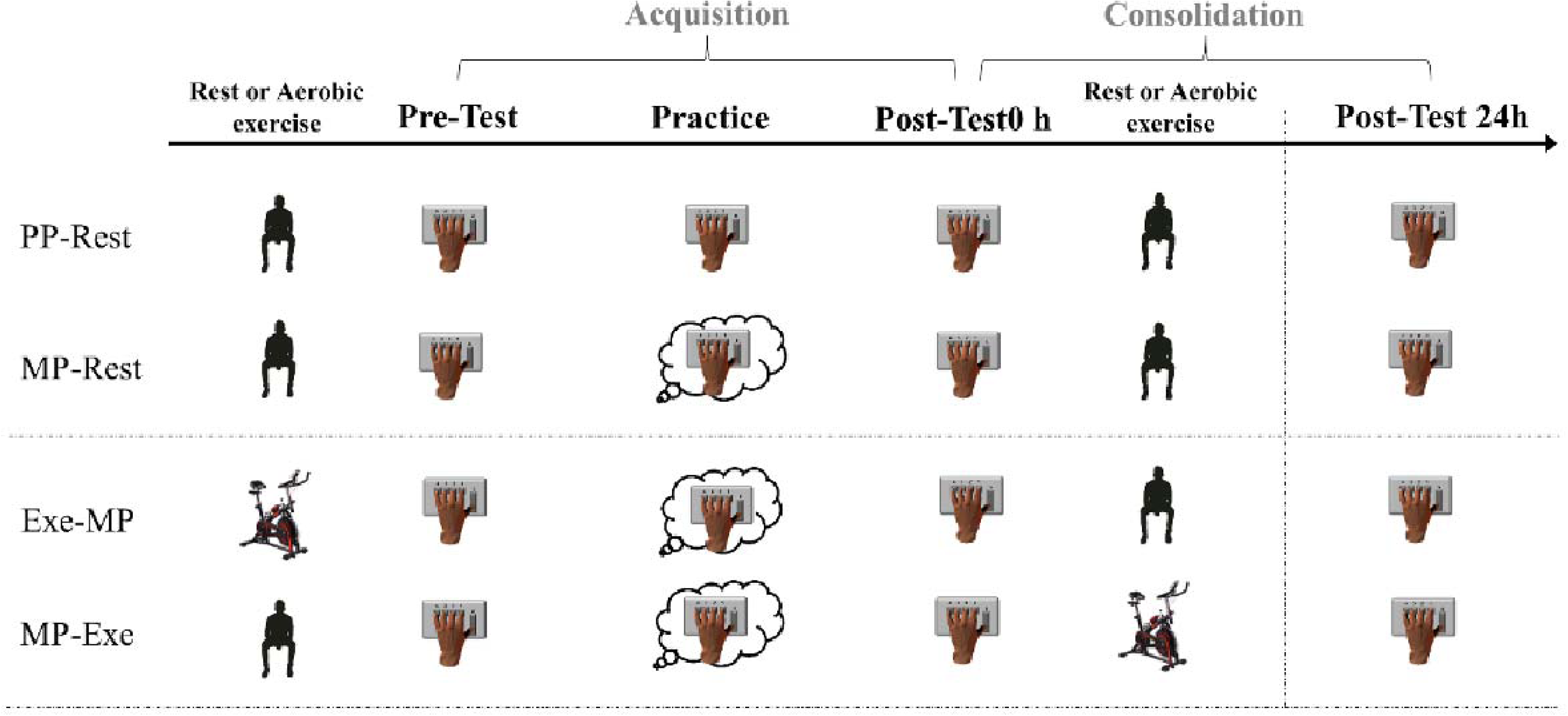
Schematic representation of the experimental procedure. Before the Pre-Test session, participants of the Exe-MP group were assigned to a moderate-intensity exercise of 30 minutes; the participants of the other groups were at rest. During the Pre-Test session, all participants actually performed two trials (30 seconds) of the sequential finger-tapping task. Participants were then assigned to a physical (PP) or a mental practice (MP) session. During the Post-Test 0h session, all participants performed again two actual trials. Depending on their group, participants were then either free to leave or were asked to perform the cycling exercise (MP-Exe group). All participants returned one day after to perform two trials of the sequential finger-tapping task (Post-Test 24h).

### Data analysis and statistics

#### Motor parameters

Two motor parameters were assessed: i) movement speed, defined as the total number of sequences performed in a 30-sec trial, and ii) accuracy, defined as the number of correct sequences performed in a 30-sec trial (Walker et al., 2003). We averaged the number of sequences from both trials for each test session.

For each parameter, we calculated the individual gain between means at Pre-Test and Post-Test 0h (motor acquisition) as follows:

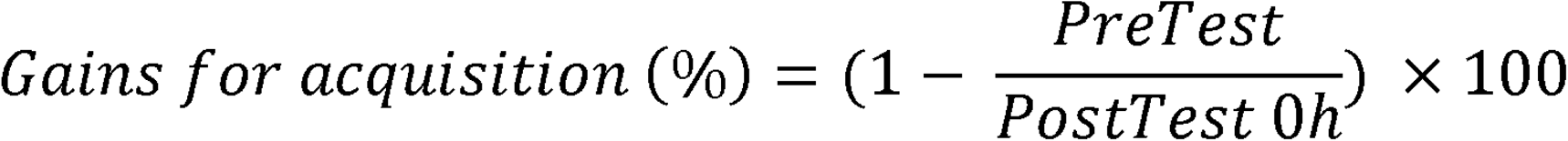

and between means at Post-Test 0h and Post-Test 24h (motor consolidation) as follows:

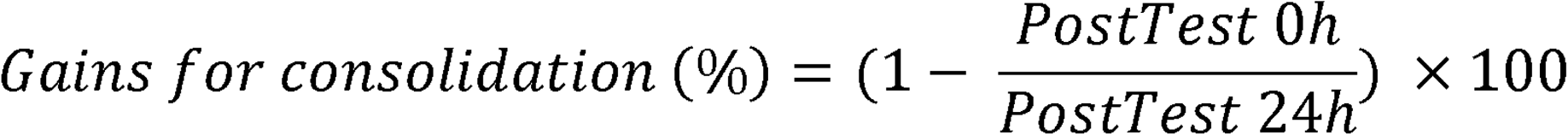

#### Electromyographic recording and analysis

Electromyographic (EMG) activity was recorded during MP to ensure the absence of muscular activity (i.e., EMG below 0.02 mV, [Mizuguchi et al., 2012]). EMG recordings were made on the left first dorsal interosseous muscle using surface Ag/AgCl electrodes in a belly-tendon montage. A ground electrode was placed on the styloid process of the ulna. The EMG signals were amplified and band-pass filtered (10–1000 Hz, Biopac Systems Inc.) and digitized at a sampling rate of 2000 Hz for offline analysis. Background EMG was monitored to ensure complete muscle relaxation throughout the experiments (EMG below 0.02 mV), using the following formula:

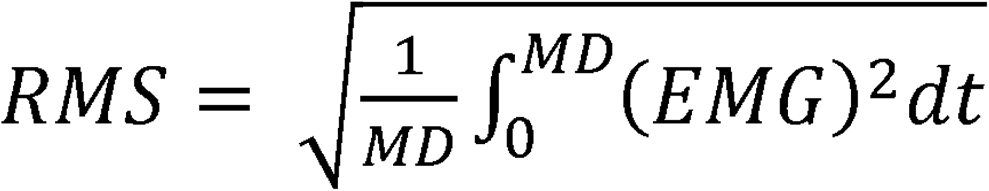

#### Statistical analysis

Statistical analyses were conducted using STATISTICA (*13.0 version; Stat-Soft, Tulsa, OK*). Normality and homogeneity of the data were tested for each parameter using the Shapiro-Wilk test and Levene test, respectively. Cohen’s d and partial eta square (η*_p_^2^*) are provided to inform on effect sizes for one-sample t-tests and ANOVAs, respectively. Pairwise comparisons (Fischer’s tests) were applied in the case of ANOVAs significance. The statistical significance threshold α was set at 0.05.

We first conducted one-factor (group) ANOVAs to control for the potential between-groups difference at Pre-Test, considering raw values for movement speed and accuracy. We also applied one-sided one-sample t-tests against the reference value 0 to test motor gains at Post-Test 0h (motor acquisition) for each parameter and each group. Holm-Bonferroni corrections were applied to adjust the P values computed for each parameter (four t-tests per parameter). Then, we ran two one-factor (Group) ANOVAs to test for between-group differences in movement speed and accuracy gains. Pairwise comparisons were managed in case of statistical significance. We performed the same analyses for motor consolidation. Finally, one-sample t-tests were conducted against the reference value of ‘0.02’ (see the “Electromyographic recording and analysis” section) to ensure EMG remained below this threshold during MP.

## Results

Table 1 summarizes the main results of the study by depicting mean values and standard deviations (SD) for the speed and accuracy parameters as well as for the different test sessions and groups. All groups showed similar performances in the Pre-Test as ANOVAs did not yield differences between groups for both movement speed (*F*_3,36_=0.93, *p*=0.43) and accuracy (*F*_3,36_=0.82, *p*=0.49).

**Table 1:** Means and standard deviations (SD) for each group for the three test sessions. PP-Rest: Physical practice without exercise; MP-Rest: Mental practice without exercise; Exe-MP: Mental practice preceded by exercise; MP-Exe: Mental practice followed by exercise; MS: Movement speed (total number of sequences); Acc: Accuracy (number of correct sequences).

|  |  | Pre-Test | Post-Test 0h | Post-Test 24h | Gains for acquisition (%) | Gains for consolidation (%) |
| --- | --- | --- | --- | --- | --- | --- |
|  |  | Mean (SD) | Mean (SD) | Mean (SD) | Mean (SD) | Mean (SD) |
| **PP-Rest** | *MS* | 12.35 (3.25) | 17.45 (4.69) | 19.9 (5.37) | 28.28 (10.10) | 12.37 (5.94) |
|  | *Acc* | 11.25 (3.49) | 15.4 (4.83) | 18.1 (4.15) | 24.81 (16.71) | 16.46 (10.40) |
| **MP-Rest** | *MS* | 15.3 (4.65) | 20.1 (5.34) | 20.45 (5.52) | 24.60 (7.91) | 0.54 (12.91) |
|  | *Acc* | 13.65 (4.23) | 18.25 (5) | 18.85 (6.42) | 25.37 (10.01) | -1.38 (19.82) |
| **Exe-MP** | *MS* | 12.65 (6.96) | 17.25 (5.89) | 17.60 (5.70) | 30.31 (16.26) | 2.21 (5.09) |
|  | *Acc* | 11.35 (7.11) | 15.4 (4.40) | 16.45 (5.25) | 30.61 (25.57) | 5.46 (9.72) |
| **MP-Exe** | *MS* | 11.25 (6.73) | 14.6 (6.29) | 15.75 (6.05) | 26.24 (15.89) | 8.94 (10.77) |
|  | *Acc* | 9.85 (6.35) | 12.95 (5.42) | 14.5 (5.07) | 28.05 (21.64) | 12.46 (11.79) |

### Gains in acquisition

Figure 2 illustrates motor gains between the Pre-Test and Post-Test 0h. One sample t-tests against the reference value “0” revealed that all groups improved their performance between Pre-Test and Post-Test 0h for both movement speed and accuracy (all *p’s*<0.05, Table 1 and Figure 2). However, ANOVAs yielded no differences between groups (Movement speed: *F*_3,36_=0.36, *p*=0.78; Accuracy: *F*_3,36_=0.26, *p*=0.86). These findings mainly suggest that moderate exercise preceding MP did not further promote motor acquisition.

**Figure 2:**
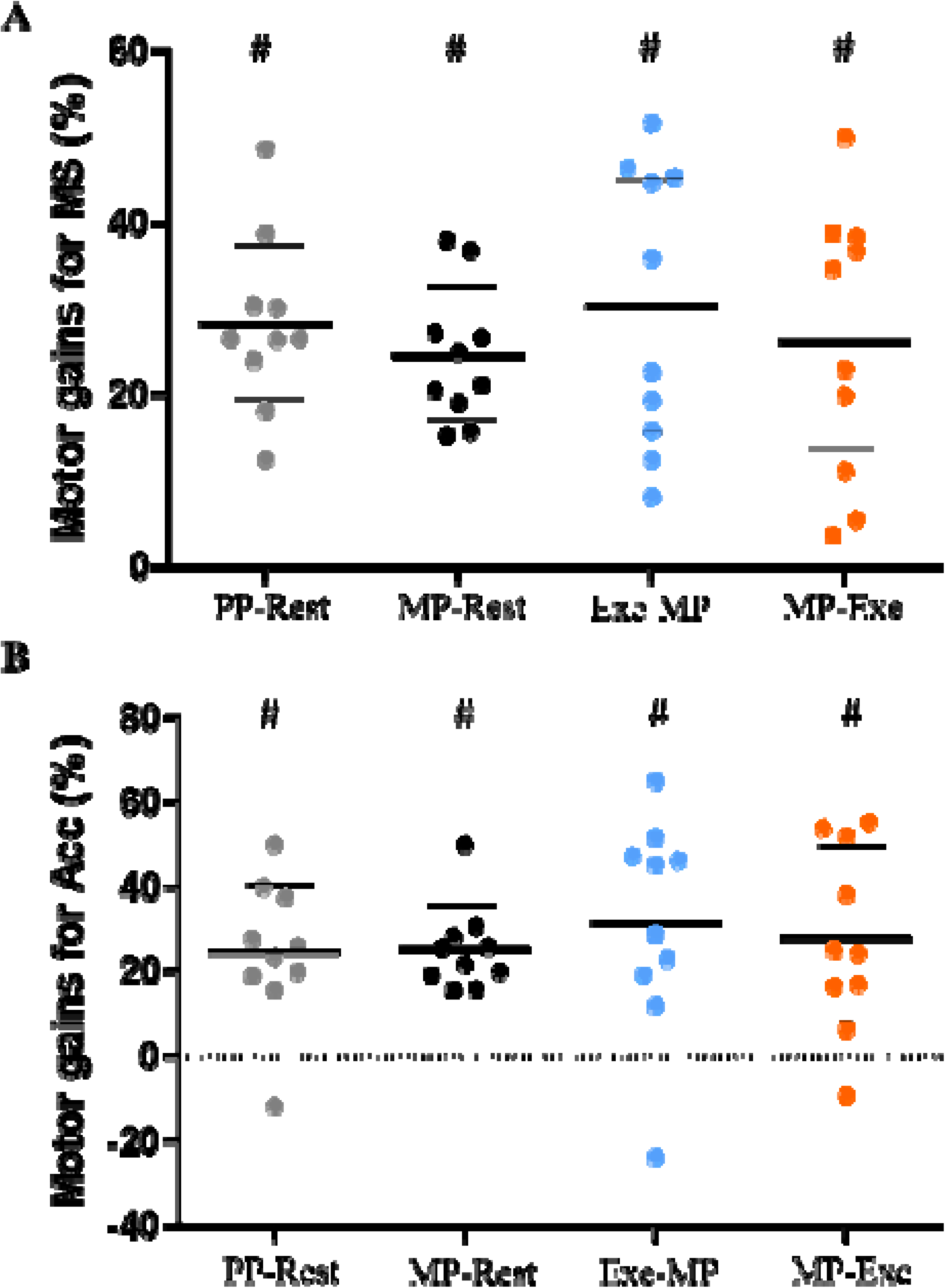
*Gains for movement speed (A) and Accuracy (B) at Post-Test 0h. Data are normalized to Pre-Test. Thick and thin horizontal lines mark mean and SD, respectively. Small circles represent individual data. PP-Rest: Physical practice without exercise; MP-Rest: Mental practice without exercise; Exe-MP: Mental practice preceded by exercise, MP-Exe: Mental Practice followed by exercise; MS: movement speed; Acc: Accuracy; #: p<0.05 (comparison to 0)*.

### Gains for consolidation

Figure 3 shows motor gains between Post-Test 0h and Post-Test 24h. One sample t-tests against the reference value “0” yielded no significant improvements (in all, *p’s*>0.05) for the MP-Rest and Exe-MP groups, suggesting a stabilization for both movement speed (MP-Rest: 0.54 ±12.91%; Exe-MP: 2.21 ±5.09%) and accuracy (MP-Rest: −1.38 ±19.82%; Exe-MP: 5.46 ±9.72%). Interestingly, both PP-Rest and MP-Exe groups improved their motor performance 24 hours following practice (off-line learning) for movement speed (PP-Rest: 12.37 ±5.94%, *t*_9_=6.58, *p*<0.01, *Cohen’s d*=2.08; MP-Exe: 8.94 ±10.77% *t*_9_=2.63, *p*=0.027, *Cohen’s d*=0.83) and accuracy (PP-Rest: 16.46 ±10.4% *t*_9_=5.00, *p*<0.01, *Cohen’s d*=1.58; MP-Exe: 12.46 ±11.79% *t*_9_=3.34, *p*<0.01, *Cohen’s d=*1.05). These results suggest that moderate-intensity exercise after MP enhanced motor consolidation.

**Figure 3:**
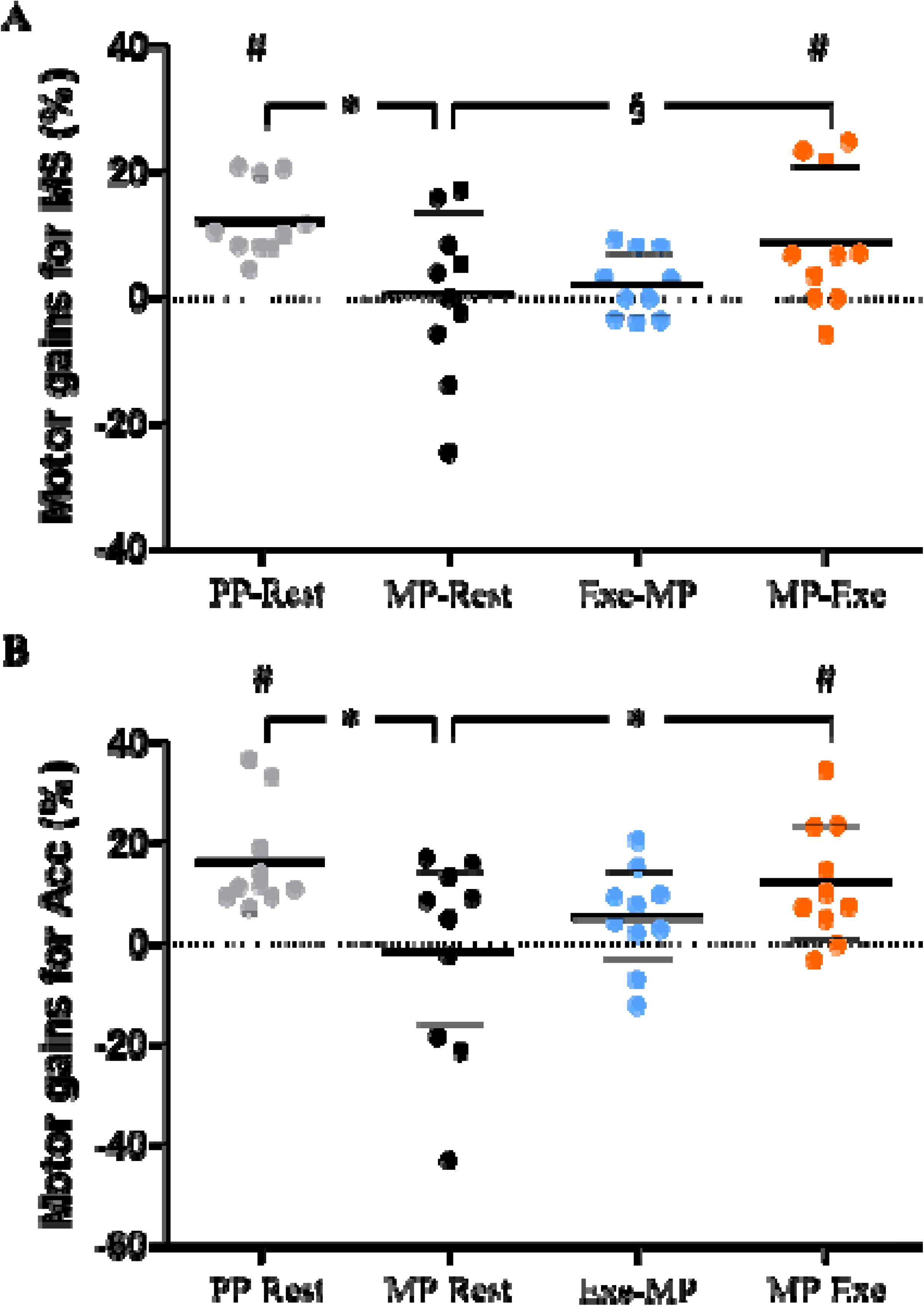
Gains for movement speed (A) and accuracy (B) at Post-test 24h. Data are normalized to Post-Test0h. Thick and thin horizontal lines mark mean and SD values, respectively. Small circles represent individual data. Brackets with asterisks refer to significant pairwise comparisons. PP-Rest: Physical practice without exercise; MP-Rest: Mental practice without exercise; Exe-MP: Mental practice preceded by exercise, MP-Exe: Mental Practice followed by exercise; MS: movement speed; Acc: Accuracy; #: p<0.05 (comparison to 0); *: p<0.05 (pairwise comparisons); §: p=0.0501 (pairwise comparison).

ANOVAs yielded a significant effect of Group for both movement speed (*F*_3,36_=3.62, *p*=0.022, η*_p_^2^*=0.23) and accuracy (*F*_3,36_=3.37, *p*=0.028, η*_p_^2^*=0.22). Considering movement speed, pairwise comparisons revealed a significant difference between the PP-Rest group and the MP-Rest group (*p*<0.01, *Cohen’s d=*1.26). The difference between the MP-Exe group and the MP-Rest group was not significant (*p*=0.0501), but with a moderate effect (*Cohen’s d=*0,71). Considering accuracy, pairwise comparisons showed significant differences between the PP-Rest and the MP-Rest groups (*p*<0.01, *Cohen’s d=*1.18) and between the MP-Exe and the MP-Rest groups (*p*=0.028, *Cohen’s d=*0.88). It is worth noting that MP-Exe and PP-Rest were statistically comparable for both movement speed (*p*=0.41, *Cohen’s d=*0,41) and accuracy (*p*=0.51, *Cohen’s d=*0,36).

Overall, the current results suggest beneficial effects of exercise following MP on motor consolidation, especially for accuracy.

### Electromyographic recordings during mental practice

EMG remained below 20 μV for each group and each block (Table 2), ensuring that participants were at rest during MP (in all, *p’s*<0.05).

**Table 2:** Means and standard deviations (SD) of electromyographic (EMG) activity of the first dorsal interosseus muscle of the left hand, recorded during mental practice (3 blocks). MP-Rest: Mental practice without exercise; Exe-MP: Mental practice preceded by exercise; MP-Exe: Mental practice followed by exercise

| **EMG activity in $\mu\text{V}$ (mean ±SD) during mental practice** |  |  |  |
| --- | --- | --- | --- |
|  | **Block 1** | **Block 2** | **Block 3** |
| **MP-rest group** | 2.06 ±1.72 | 1.93 ±1.96 | 1.91 ±1.72 |
| **Exe-MP group** | 1.43 ±0.66 | 1.28 ±0.58 | 1.30 ±0.42 |
| **MP-Exe group** | 1.26 ±1.26 | 1.02 ±1.02 | 1.26 ±1.06 |

## Discussion

In this study, we investigated the effect of moderate-intensity exercise on motor acquisition and consolidation induced by MP. Consistently with previous reports, we showed a significant improvement in motor performance, that is in movement speed and accuracy, immediately after practice for both MP and PP. In contradiction with our first hypothesis, moderate-intensity exercise potentiated neither motor acquisition nor motor consolidation when preceding MP. Interestingly, when following MP, moderate-intensity exercise further improved motor consolidation, supporting thus the second hypothesis. The performance was at the level of the PP group that only performed actual trials during practice (i.e., without extra exercise).

### Improvement in motor performance after PP and MP

The current findings are in accordance with the previous literature, showing that both MP and PP lead to the accomplishment of faster and more accurate movements (Gentili et al., 2006, 2010; Rannaud Monany et al., 2022a). It is currently assumed that the improvement of performance induced by MP and PP relies on neural adaptations that occur through an overlapping cortico-subcortical network. For example, previous work by Avanzino et al. (2015) and Bonassi et al. (2017) showed that both MP and PP induce LTP-like plasticity and increase corticospinal excitability at the level of the primary motor cortex after motor acquisition. Also, Pascual-Leone et al. (1995) suggested that extended MP and PP both lead to motor consolidation through cortical reorganization mechanisms. This is supported by more recent of Grosprêtre et al. (2017), who showed that extended mental practice (1 week) increased excitability at both cortical and spinal levels. It is worth noting that neural adaptations induced by MP and PP have also been reported at the level of other sensorimotor areas, notably including premotor areas and the cerebellum (Lacourse et al., 2004). Altogether, these studies support the involvement of shared plasticity mechanisms for MP and PP concerning motor acquisition and consolidation.

### The non-significant effect of moderate-intensity exercise on motor acquisition

As a reminder, we hypothesized that moderate-intensity exercise before MP would promote motor acquisition and increase motor performance compared to MP without exercise. This hypothesis was raised based on previous work of Statton et al. (2015) on PP. Specifically, these authors observed that moderate-intensity exercise before PP promoted motor acquisition and led to better motor performance when compared to PP without exercise. It is proposed that moderate-intensity exercise enhances motor acquisition by acting on psychological and neuroendocrinological factors, as well as neuroplastic mechanisms. Previous work showed, for example, that moderate-intensity exercise increases arousal, which has been associated with better cognitive performances (Lambourne and Temporoski, 2010). Exercise also modulates the circulation of various neuroendocrine substances, such as dopamine, that are implied in motor acquisition (Foley and Fleshner, 2008; de Sousa Fernandes et al., 2020). Importantly, moderate-intensity exercise facilitates the induction of LTP-like plasticity during PP by transiently reducing short-interval intracortical inhibition (SICI) at the level of the primary motor cortex (Devanne and Allart, 2019; Singh et al., 2014, 2015). Considering this mechanism, we suggest that the absence of benefits of moderate-intensity exercise before MP may partly rely on the effects of motor imagery on SICI. Indeed, it is known that motor imagery involves interactions between excitatory and inhibitory processes at the level of the primary motor cortex (Grosprêtre et al., 2017). Specifically, recent work of our team (Neige et al., 2020) pointed out that SICI is likely to increase during motor imagery, perhaps to prevent movement execution while imagining. Accordingly, we consider that there may be a conflict between the lowering and increasing of SICI, respectively due to moderate-intensity exercise and motor imagery. The increase of SICI during motor imagery could have interfered with the effects of moderate-intensity exercise on LTP-like plasticity induction during MP and thus reduced its benefits on motor acquisition.

### Moderate-intensity exercise after MP potentiates motor consolidation

The current results revealed a stabilization (MP-Rest and Exe-MP) or an improvement (PP-Rest and MP-Exe) of motor performance between Post-Test 0h and Post-Test 24h, attesting a motor consolidation (Krakauer and Shadmehr, 2006; Walker et al., 2003). Interestingly, moderate-intensity exercise after MP potentiated motor consolidation, leading to better motor performances when compared to MP without exercise. The ways that moderate-intensity exercise potentiates motor consolidation are numerous and not fully identified, even for PP. Previous reports present the increase of lactate and BDNF levels induced by exercise as important factors to explain the benefits of motor consolidation (Skriver et al., 2014). Nonetheless, the effects of moderate-intensity exercise on these biomarkers are intensity-dependent, meaning that high-intensity exercises are more likely to induce such increase than moderate- or low-intensities ones (Ferris et al., 2007). Reports are less consistent in the case of moderate-intensity exercise. One study however pointed out that a bout of moderate-intensity exercise increased the activity of the neuro-adrenergic system (Segal et al., 2012), which is involved in motor consolidation (Kuo et al., 2021).

To our knowledge, only one study investigated the effects of moderate-intensity exercise on motor consolidation when performed after PP (Thomas et al., 2016). In that study, authors found positive effects of exercise on motor consolidation seven days after PP but not one day after. Relating to current findings, an increase in motor performance questions the potential differences between MP and PP on motor consolidation. Recent behavioral studies suggested that motor consolidation after MP and PP may rely on distinct processes (Ruffino et al., 2021). This is corroborated by neurophysiological experiments which suggest that motor consolidation by MP and PP leads to different patterns of brain activity (Kraeutner et al., 2020). Also, and according to Truong et al. (2022), motor consolidation by MP would involve slower processes than PP, which notably intervene during the passage of time that follows motor acquisition. In agreement with that hypothesis, we speculate that moderate-intensity exercise could have promoted such processes when performed after MP, explaining the better consolidation for the MP-Exe group when compared to the MP-Rest group. Further investigations will be required to elucidate the neural processes that sustain motor consolidation following MP, as well as the effects of moderate-intensity exercise on these processes.

The current work contains some limitations. It is worth noting that the implementation of neuromodulation protocols, such as transcranial magnetic stimulation, to measure corticospinal excitability and SICI, would have been highly beneficial to support our interpretations. Also, measurements of perceived exercise intensity, using effort perception scales, would have been of interest to ensure that exercise was perceived as moderate by the participants.

## Perspectives

The current results support the benefits of moderate-intensity exercise on motor consolidation following MP with motor imagery. While MP has been shown to improve motor performance overnight (Freitas et al., 2020; Truong et al., 2022), optimizing motor imagery-based protocols are of importance for sports medicine (Rannaud Monany, 2022b). The intensity of exercise would be a key point to induce plasticity mechanisms and to promote motor consolidation following MP.

## Author contributions

DRM, CP, and FL designed the experiment, DRM conducted the experiment and prepared the figures, DRM and FL analyzed the data, and DRM, CP, and FL wrote the manuscript.

## Data Availability Statement

The data are available at https://osf.io/4zyud

